# Mitochondrial membrane potential acts as a retrograde signal to regulate cell cycle progression

**DOI:** 10.1101/2022.02.18.480979

**Authors:** Choco Michael Gorospe, Alicia Herrera Curbelo, Gustavo Carvalho, Lisa Marchhart, Katarzyna Niedźwiecka, Paulina H. Wanrooij

## Abstract

Mitochondria are central to numerous anabolic and catabolic pathways whereby mitochondrial dysfunction has a profound impact on metabolism and can manifest in disease. The consequences of mitochondrial dysfunction can be ameliorated by adaptive responses that rely on mito-cellular crosstalk to communicate mitochondrial distress to the rest of the cell. Such mito-cellular signaling slows cell cycle progression in mitochondrial-DNA deficient (ρ^0^) *Saccharomyces cerevisiae* cells, but the initial trigger and the pathway mediating the response has remained unknown. Here, we show that decreased mitochondrial membrane potential (ΔΨm) acts as the initial signal of mitochondrial stress that delays G1-to-S phase transition in both ρ^0^ and control cells. Accordingly, experimentally increasing ΔΨm was sufficient to restore timely cell cycle progression in ρ^0^ cells. Neither the RTG retrograde pathway nor central DNA damage checkpoint kinases were involved in mediating this form of mito-cellular communication. The identification of ΔΨm as a novel regulator of cell cycle progression may have implications for disease states involving mitochondrial dysfunction.

## Introduction

Mitochondria use the electron transport chain (ETC) complexes to convert energy gained from the oxidation of nutrients into an electrochemical gradient across the inner mitochondrial membrane; this electrochemical gradient is then used to drive ATP synthesis through the process of oxidative phosphorylation (OXPHOS). In addition to producing the majority of the cell’s ATP, however, mitochondria carry out a diverse array of vital cellular functions including synthesis of Fe-S clusters, amino acid and nucleotide biosynthesis, the production of reactive oxygen species (ROS) and apoptosis. It follows that mitochondrial function is required for cell survival even in a facultative anaerobe like the budding yeast *Saccharomyces cerevisiae* that can survive without mitochondrial ATP production. In line with the manifold functions of the organelle, mitochondrial dysfunction is associated with numerous diseases including neurodegenerative disorders, metabolic syndrome, cancer, as well as aging (Nunnari and Suomalainen, 2012; Raimundo, 2014). Rather than manifesting merely as an energy defect, mitochondrial dysfunction can contribute to cellular dysfunction and disease etiology through diverse mechanisms involving *e*.*g*. increased levels of ROS that can damage cellular constituents, changes in nuclear epigenetic marks or gene expression patterns, and even instability of the nuclear genome (Leadsham et al., 2013; Nunnari and Suomalainen, 2012; Kopinski et al., 2019; Veatch et al., 2009).

To avoid or ameliorate the dire consequences of mitochondrial dysfunction, a complex communication network mediates signals of mitochondrial status to other parts of the cell including the nucleus, lysosomes, peroxisomes and the endoplasmic reticulum (Nunnari and Suomalainen, 2012; Raimundo, 2014). This mito-cellular signaling can be mediated by signals ranging from key metabolites to misfolded proteins or ROS, and acts to restore cellular homeostasis, facilitate adaptation to the altered mitochondrial status, or eliminate dysfunctional mitochondria via mitophagy (Mottis et al., 2019).

The most thoroughly understood pathway of mito-cellular communication is the RTG (<u>r</u>e<u>t</u>ro<u>g</u>rade) pathway in *S. cerevisiae* that is activated in response to mitochondrial dysfunction such as respiratory chain deficiency or loss of mitochondrial DNA (mtDNA) (Parikh et al., 1987; Liu and Butow, 2006). The RTG pathway relies on the heterodimeric Rtg1-Rtg3 transcription factor to activate the expression of RTG target genes and ensure sufficient synthesis of glutamate – the main source of nitrogen in biosynthetic processes – even in the absence of mitochondrial function (Liu and Butow, 2006). Activation of the RTG response requires the positive regulator Rtg2 that promotes the partial dephosphorylation and consequent nuclear translocation of the normally cytosolic Rtg1-Rtg3 complex, thus enabling activation of the RTG response. The initial signal triggering this retrograde pathway has been reported to be the drop in mitochondrial membrane potential (ΔΨm) that accompanies loss of mtDNA and respiratory deficiency (Miceli et al., 2012).

MtDNA encodes key subunits of the ETC complexes as well as the F_o_ component of the ATP synthase, making this small genome essential for cellular respiration and OXPHOS. It follows that cells lacking mtDNA (termed ρ^0^ cells) cannot maintain ΔΨm by proton pumping; instead, they maintain a limited ΔΨm by a mechanism involving the hydrolysis of ATP by the “reverse” action of the F_1_ component of the ATP synthase in consort with the electrogenic exchange of ATP^4-^ for ADP^3-^ by the adenine nucleotide translocator ANT (Giraud and Velours, 1997; Buchet and Godinot, 1998; Appleby et al., 1999). Certain mutations in subunits of the F_1_-ATPase increase the activity of the enzyme, resulting in a higher ANT-driven membrane potential in ρ^0^ cells (Kominsky et al., 2002).

In addition to preventing cellular respiration and decreasing ΔΨm, mtDNA loss promotes nuclear DNA instability and defective cell cycle progression that manifests as an accumulation of cells in G1 phase (Veatch et al., 2009; Zyrina et al., 2015). In an elegant study, Veatch *et al*. showed these two effects to be separate from each other: while the nuclear DNA instability in mtDNA-depleted cells was driven by defective Fe-S cluster protein assembly, the mechanism(s) underlying the cell cycle defect were not uncovered but were found to be independent of Fe-S metabolism (Veatch et al., 2009). A subsequent study suggested the cell cycle phenotype of ρ^0^ cells – termed the mtDNA inheritance checkpoint – to be dependent on the Rad53 checkpoint kinase that is central in the cell’s response to nuclear DNA damage or replication stress (Crider et al., 2012). In canonical checkpoint signaling, Rad53 functions downstream of the Mec1 master kinase that is activated in response to stretches of Replication Protein A-coated single-stranded DNA, a sign of replication stress or DNA damage (Wanrooij et al., 2016; Navadgi Patil and Burgers, 2011). Once active, Mec1 stimulates Rad53 to phosphorylate numerous downstream targets to arrest cell cycle progression and upregulate DNA repair in order to ensure the damage is resolved prior to cell division. A second master kinase of the same phosphoinositide 3-kinase related kinase family as Mec1, Tel1, phosphorylates a partly-overlapping network of targets to guarantee the timely repair of double-stranded DNA breaks (Harrison and Haber, 2006). The involvement of the Mec1 and Tel1 master kinases in the cell cycle phenotype of ρ^0^ cells has to our knowledge not been addressed.

Despite the undisputed involvement of mitochondrial dysfunction in human disease, the molecular-level events that link mtDNA loss to the cell cycle machinery, as well as the mitochondria-proximal signal that initiates this form of mito-cellular communication remain uncharacterized. Here, we analyzed the dependence of the ρ^0^ cell cycle defect on canonical checkpoint proteins and on the RTG signaling pathway, and find both to be dispensable for the G1-to-S phase progression delay known as the mtDNA inheritance checkpoint. Remarkably, a combination of genetic and pharmaceutical interventions uncovered decreased mitochondrial membrane potential as a regulator of cell cycle progression in both ρ^+^ and ρ^0^ cells, while inhibition of mitochondrial ATP synthesis or altered ROS levels did not account for the cell cycle phenotype of mtDNA-depleted cells. Finally, we show that the cell cycle defect in ρ^0^ cells can be rescued by increasing ΔΨm, confirming that adequate ΔΨm rather than mitochondrial function *per se* is required for normal cell cycle progression. These findings identify the mitochondrial membrane potential as a novel regulator of cell cycle progression.

## Results

### P^0^ cells exhibit slow growth characterized by G1-to-S phase transition delay

In order to study the effects of mtDNA loss, an mtDNA-devoid (ρ^0^) variant of the mtDNA-containing (ρ^+^) wildtype (WT) strain was made by standard treatment with ethidium bromide (Goldring et al., 1970), and confirmed by quantitative real-time PCR. P^0^ yeast cells are respiratory-deficient and can thus only grow on fermentable carbon sources such as dextrose. In accordance with previous reports, ρ^0^ cells formed characteristic small colonies on dextrose-containing rich medium, and grew slower than their ρ^+^ counterparts, unable to exceed an OD_600_ of *ca* 6 even after 48 h in liquid culture (Fig. 1A; Fig. S1A) (Ephrussi, 1949). Furthermore, the percentage of G1 cells in an early-logarithmic unsynchronized culture of ρ^0^ cells was twice as high as in ρ^+^ cells grown under identical conditions; this difference remained constant for at least 2 h during logarithmic growth (Fig. 1B-C; Fig. S1B). To test whether the increased frequency of G1 cells in the unsynchronized ρ^0^ culture was due to a slower G1-to-S transition or a faster progression through other cell cycle stages, we synchronized cells at the G1/S boundary using the α-factor pheromone and compared the timing of S phase entry in ρ^+^ and ρ^0^ cells. Following release from G1 synchrony, the number of ρ^+^ cells in G1 phase started to decrease after *ca* 20 min, signifying entry into S phase (Fig. 1D-E). However, ρ^0^ cells took significantly longer to transition from G1 to S phase, as the frequency of G1 cells did not begin to decrease until over 30 min after release from α-factor synchrony. To confirm that the G1-to-S transition delay was a true phenotype of the ρ^0^ variant and not an artifact of the α-factor treatment or an overall slower transition through the cell cycle, cells were next synchronized in G2 by nocodazole treatment. Interestingly, ρ^+^ and ρ^0^ cells released from nocodazole synchrony showed comparable timing of G2 exit and entered the G1 phase around 50 minutes after release (Fig. 1F; Fig. S1C-D). However, while ρ^+^ cells continued straight into S phase (as shown by an accumulation of S phase cells peaking at 70 min), ρ^0^ cells accumulated in G1 and displayed delayed entry into S phase (Fig. 1F). Consequenty, the percentage of ρ^0^ cells in S phase did not peak until 90-100 min after release from G2, 20-30 min later than ρ^+^ cells (Fig. 1F).

**Fig. 1.**
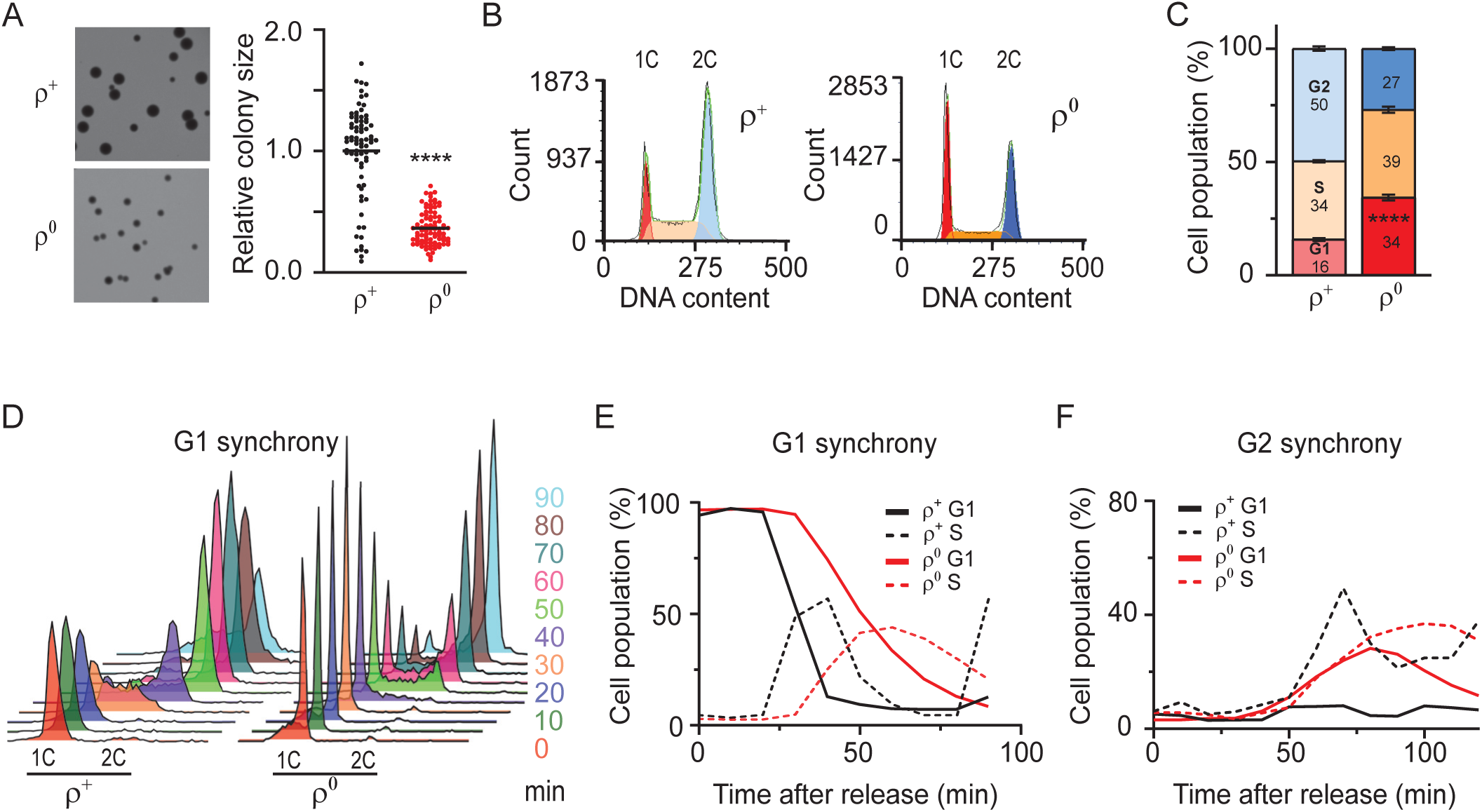
Loss of mtDNA induces a delay in transition from G1 to S phase of the cell cycle. **(A)** Growth of wild-type (WT; AC403) ρ^+^ and ρ^0^ cells in YPDA solid medium imaged after 48 h (*left panel*), and colony area measured relative to WT ρ^+^ (*right panel*). **(B)** Representative DNA histogram of unsynchronized WT (AC402) ρ^+^ and ρ^0^ cells grown to early logarithmic phase in YPDA. *1C* and *2C* indicate populations with single and double chromosome content, corresponding to cells in G1 and G2, respectively. The cell cycle profile was analyzed using the multicycle model in FCS Express to determine the percentage of cells in G1 (red), S (orange) and G2 (blue) phase. **(C)** Quantification of the percentage of cells in G1, S or G2 phase. Values represent the average of at least 4 independent experiments including the one in Fig. 1B, and error bars indicate standard deviation. The two-tailed Student’s t-test was performed to determine statistical significance between the G1 populations in WT ρ^+^ and ρ^0^ cultures. ****p <0.0001. **(D)** DNA histogram of WT (AC402) ρ^+^ and ρ^0^ cells released from G1 synchrony achieved by treatment with 10 μg/ml α-factor. Cells were sampled every 10 minutes after release. The experiment was repeated at least three times; a representative experiment is shown. **(E)** Quantification of the percentage of G1 (*solid lines*) and S phase (*dashed lines*) cells after release from G1 synchrony in the experiment shown in Fig. 1D. **(F)** Quantification of the percentage of G1 (*solid lines*) and S phase (*dashed lines*) cells after release of WT ρ^+^ and ρ^0^ cells from G2 synchrony achieved by nocodazole treatment. The DNA histograms are shown in Fig. S1B. Values represent data from a single experiment. See also Fig. S1.

Together, these results demonstrate that ρ^0^ cells exhibit slow growth characterized by a delayed G1-to-S transition, and thus confirm previous observations of an impaired G1 exit upon mtDNA loss that has been termed the mtDNA inheritance checkpoint (Crider et al., 2012; Veatch et al., 2009; Zyrina et al., 2015). However, the factors mediating the mtDNA inheritance checkpoint, as well as the initial signal that triggers the cell cycle defect, have not been uncovered. Given that this checkpoint has been suggested to be dependent on the Rad53 checkpoint kinase (Crider et al., 2012), we first investigated the requirement of central checkpoint factors for the G1-to-S delay in ρ^0^ cells.

### The G1-S delay in ρ^0^ cells is not dependent on central checkpoint kinases including Rad53

The canonical checkpoint responses that are triggered upon nuclear DNA damage or replication defects are driven by the master checkpoint kinases Tel1 and Mec1, the orthologs of mammalian ATM and ATR, respectively (Beyer and Weinert, 2014). Mec1 has also been reported to be required for efficient G1 arrest upon nutrient deprivation, a response that could be activated in ρ^0^ cells (Weinberger et al., 2007). We therefore tested the contribution of Tel1 and Mec1 to the G1-to-S delay in ρ^0^ cells by assessing the cell cycle profile of the relevant deletion mutants and their ρ^0^ derivatives. Interestingly, the deletion of *TEL1* did not prevent the accumulation of ρ^0^ cells in G1 phase, indicating that the G1-to-S delay observed in ρ^0^ cells is independent of the Tel1 checkpoint kinase (Fig. 2A; Fig. S2A). We next tested the involvement of Mec1. As null mutation of *MEC1* is lethal unless accompanied by deletion of the gene encoding the ribonucleotide reductase inhibitor Sml1, the analyses were carried out in a *sml1*Δ background (Zhao et al., 1998). The cell cycle profiles of the *sml1*Δ ρ^+^ and ρ^0^ cells were comparable to their WT counterparts (Fig. 2B; Fig. S2A). Similar to *TEL1*, deletion of *MEC1* in *sml1*Δ cells did not prevent the increase in G1 cells in the ρ^0^ variant (Fig. 2B; Fig. S2A). Furthermore, and in accordance with Fig. 2A, deletion of *TEL1* in *sml1*Δ cells did not alter the cell cycle profile of ρ^+^ or ρ^0^ cells (Fig. S2B). We conclude that neither the Tel1 nor the Mec1 master checkpoint kinase are required for the delayed G1-to-S transition observed in ρ^0^ cells.

**Fig. 2.**
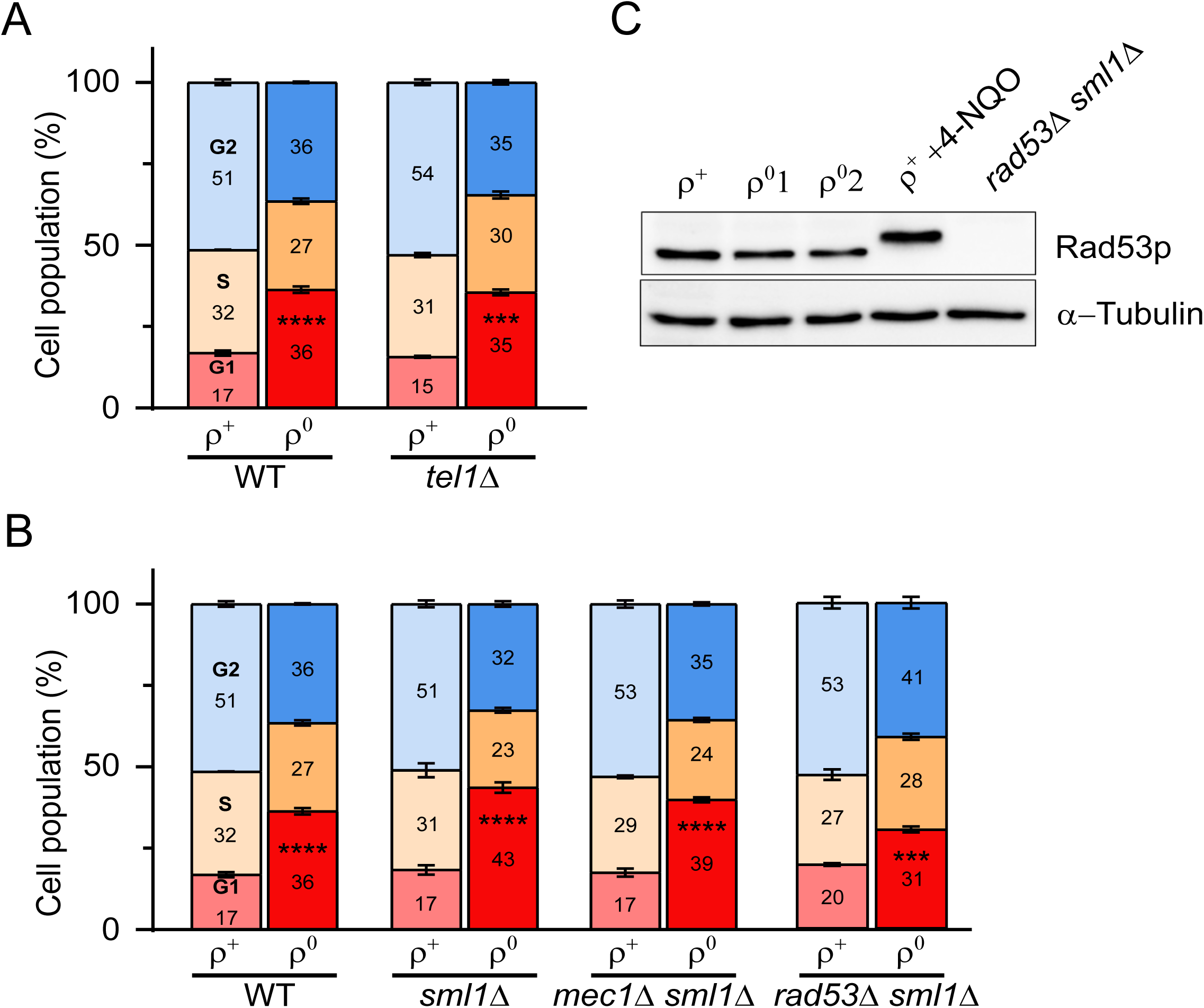
The Tel1, Mec1 and Rad53 checkpoint kinases are redundant for the G1-S delay in ρ^0^ cells. **(A-B)** Quantification of the cell cycle profiles of WT, *tel1*Δ, *sml1*Δ, *mec1*Δ *sml1*Δ and *rad53*Δ *sml1*Δ ρ^+^ and ρ^0^ cells. Values represent the average of at least 3 independent experiments and error bars indicate standard deviation. The G1 population in each ρ^+^ and its ρ^0^ variant was compared using the two-tailed Student’s t-test. ****p<0.0001, ***p<0.001. The same WT data is presented in both panels. Representative DNA histograms are shown in Fig. S2A. **(C)** Western blot analysis of Rad53 and α-Tubulin in WT ρ^+^ and ρ^0^ cells harvested in early log phase. WT ρ^+^ cells treated with 4-NQO for 60 min before harvesting served as a positive control for phosphorylation and the consequent shift in migration of Rad53p. *rad53*Δ *sml1*Δ ρ^+^ cells served as the negative control. See also Fig. S2.

In canonical checkpoint signaling, activated Mec1 phosphorylates and activates the downstream Rad53 effector kinase (Sanchez et al., 1996; Hoch et al., 2013). Given that we found the G1-to-S transition delay in ρ^0^ cells to be independent of Mec1, we revisited the suggested dependence of this phenotype on Rad53. Similar to *MEC1, RAD53* is an essential gene and its deletion can only be achieved via prior inactivation of the *SML1* gene. The percentage of G1 and S phase cells in *rad53*Δ *sml1*Δ ρ^+^ cultures did not significantly differ from *sml1*Δ ρ^+^ cells (Fig. 2B; Fig. S2A). Similarly, *rad53*Δ *sml1*Δ cells showed an increased percentage of G1 cells in ρ^0^ compared to ρ^+^ cells, although the increase was somewhat more modest than in *sml1*Δ (G1 cells in *rad53*Δ *sml1*Δ increased from 20 to 31% after conversion from ρ^+^ to ρ^0^, respectively, compared to 17 and 43% in *sml1*Δ). These findings indicate that Rad53 is not required for the G1-to-S transition delay observed in ρ^0^ cells under our experimental conditions. We observed comparable results in another independent *rad53*Δ *sml1*Δ strain as well as in a *rad53*Δ *sml1-1* strain in another W303-derivative background (Fig. S2C). Finally, we used Western blot analysis to directly assay for potential Rad53 activation in ρ^0^ cells. Although the gel migration of Rad53 was clearly retarded in ρ^+^ cells treated with the DNA damaging agent 4-NQO, indicating that Rad53 phosphorylation was readily detected in our assay, no phosphorylation of Rad53 was detected in WT ρ^0^ cells (Fig. 2C).

These results therefore indicate that the G1-to-S phase progression delay observed in WT ρ^0^ cells is independent of the Mec1/Rad53 and Tel1 checkpoint signaling pathways.

### The RTG-dependent retrograde signaling pathway is not required for the G1-to-S phase delay in ρ^0^ cells

The RTG pathway is the most thoroughly-understood pathway for mitochondria-to-nucleus signaling upon mitochondrial dysfunction. This retrograde pathway is activated upon mtDNA loss, and is thereby a prime candidate for mediating the signaling that leads to the G1- to-S transition delay in mtDNA-depleted cells (Liao and Butow, 1993; Kirchman et al., 1999). We first tested the effect of deleting the gene encoding the proximal sensor of the RTG pathway, *RTG2*. The cell cycle profile of *rtg2*Δ ρ^+^ and ρ^0^ cells was similar to that of their WT counterparts, and the percentage of G1 cells was considerably higher in *rtg2*Δ ρ^0^ than *rtg2*Δ ρ^+^ cells, indicating that Rtg2 was not required for the G1-to-S delay (Fig. 3 and Fig. S3). Even Rtg3, which contains the transcriptional activation domain in the Rtg1/Rtg3 complex (Rothermel et al., 1997), was found to be redundant for the G1-to-S phase delay in ρ^0^ cells (Fig. 3 and Fig. S3). We conclude that although the RTG pathway generally responds to mitochondrial distress and loss of mtDNA, it is not required for the G1-to-S phase progression delay observed in ρ^0^ cells.

**Fig. 3.**
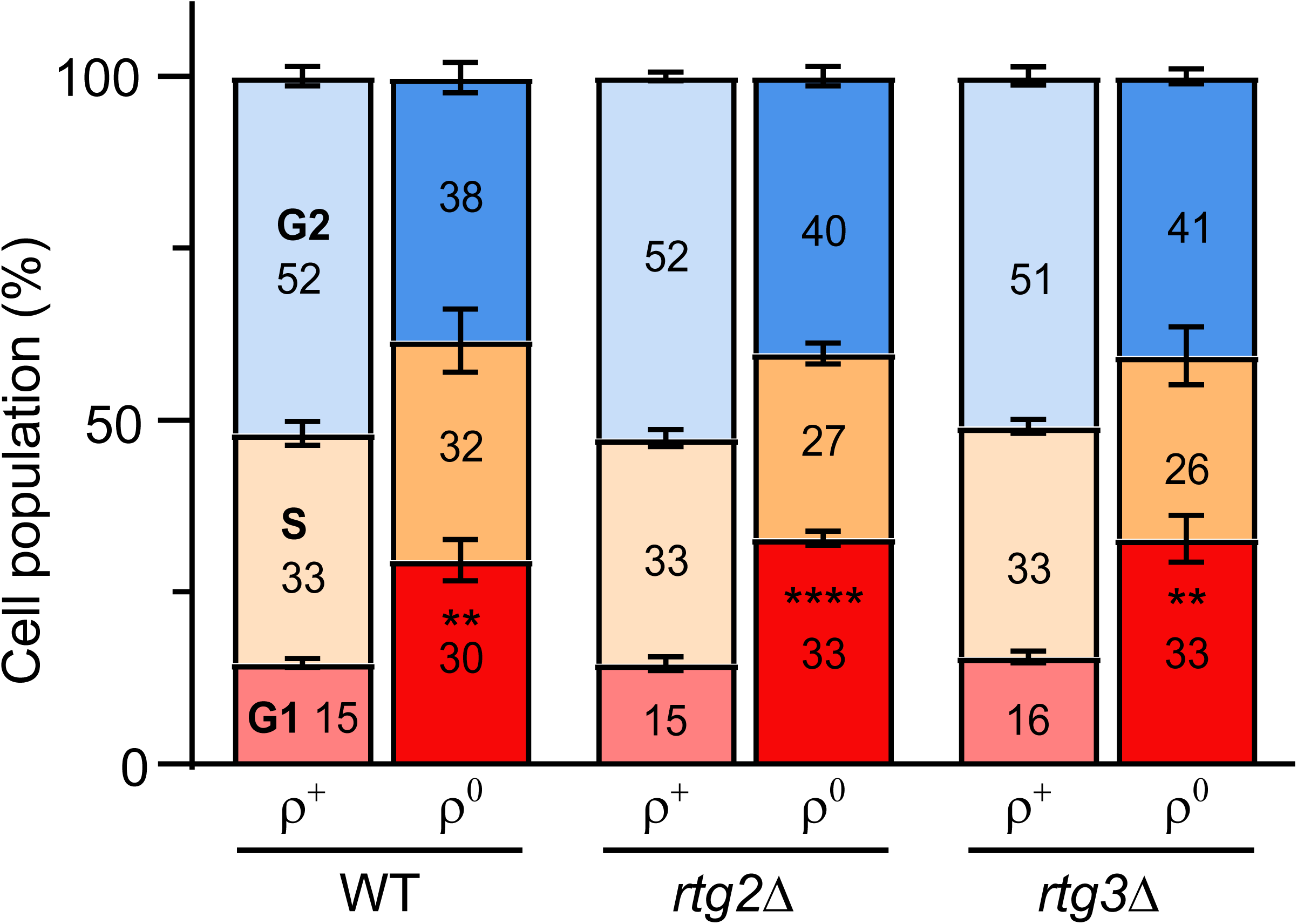
The G1-S transition delay in ρ^0^ cells does not require the RTG retrograde pathway. Quantification of the cell cycle profiles of WT, *rtg2*Δ and *rtg3*Δ ρ^+^ and ρ^0^ cells. Values represent the average of at least 4 independent experiments, and error bars indicate standard deviation. The G1 population in each ρ^+^ and its ρ^0^ variant was compared using the two-tailed Student’s t-test. ****p <0.0001, **p< 0.01. Representative DNA histograms are shown in Fig. S3.

### Decreased mitochondrial membrane potential delays G1-to-S phase progression

We next sought to identify the proximal signal that triggers the cell cycle phenotype in mtDNA-deficient cells. Yeast mtDNA encodes subunits of the ETC and the ATP synthase. Consequently, electron transport and OXPHOS are impaired in ρ^0^ cells, resulting in a plethora of outcomes including impaired mitochondrial ATP production, decreased ΔΨm, and increased generation of ROS (Greaves et al., 2012; Houten et al., 2016; Dirick et al., 2014). We reasoned that any of these downstream effects of a malfunctional ETC and OXPHOS were reasonable candidates for the initial signal that triggers the G1-to-S phase delay in ρ^0^ cells, and tested each of them in turn.

We first assessed the effect of pharmacologically inhibiting mitochondrial ATP production using oligomycin, a compound that binds to the Fo subunit of the ATP synthase to prevent the flow of protons back into the mitochondrial matrix (Lee and O’Brien, 2010). Treatment of ρ^+^ or ρ^0^ cells with oligomycin did not cause accumulation of cells in G1 phase or decrease the percentage of cells in S phase compared to untreated cells over the observation period of 120 min following treatment (Fig. 4A, Fig. S4A-B). Therefore, the G1-to-S transition delay in ρ^0^ cells is not triggered by decreased mitochondrial ATP synthesis, a conclusion that is in agreement with previous findings and the fact that we assay for the cell cycle delay during early logarithmic growth on dextrose-containing media when ATP generation primarily occurs by fermentation rather than by mitochondrial respiration (van Dijken et al., 1993; Crider et al., 2012).

**Fig. 4.**
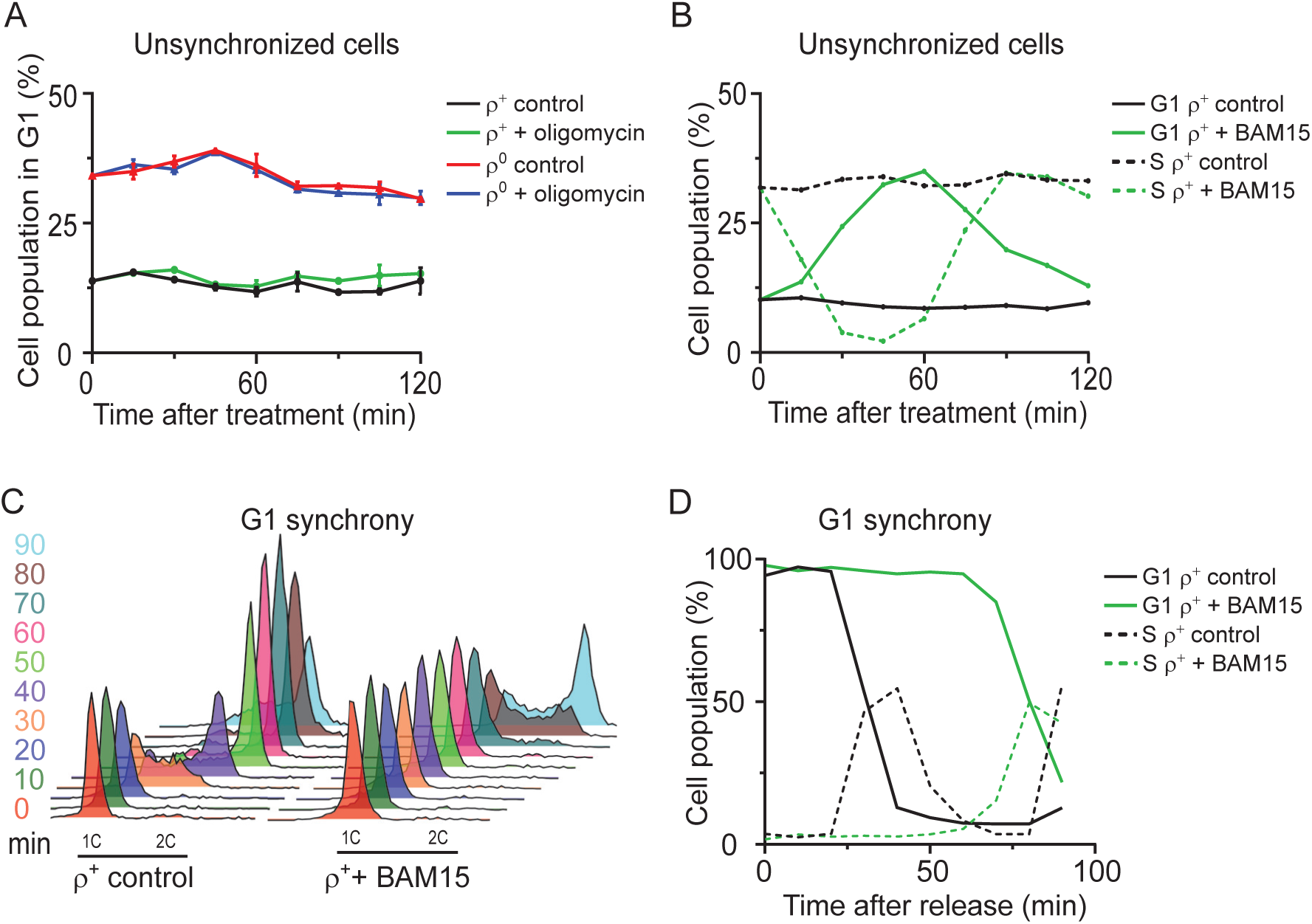
The loss of ΔΨm, but not the inhibition of mitochondrial ATP synthesis, delays G1- to-S progression in WT ρ^+^ cells. **(A)** The percentage of G1 cells in WT ρ^+^ and ρ^0^ cells left untreated or treated with 20 μM of oligomycin to inhibit mitochondrial ATP synthesis. Aliquots were harvested upon addition of the drug (0 min) and every 15 min thereafter. Values represent the average of two independent experiments, and error bars indicate standard deviation. Representative DNA histograms and the quantification of S phase cells are shown in Fig. S4A-B. **(B)** The percentage of G1 (*solid lines*) and S phase (*dashed lines*) cells in cultures of WT ρ^+^ cells left untreated or treated with 30 μM of BAM15. Aliquots were harvested upon addition of the drug (0 min) and every 15 min thereafter. Representative DNA histograms are shown in Fig. S4C. Values represent data from a single experiment repeated at least three times. **(C)** DNA histogram of WT ρ^+^ cells synchronized in G1 with 10 μg/ml α-factor and released into media with or without 30 μM BAM15. Cells were sampled upon release (time 0) and every 10 minutes thereafter. **(D)** Quantification of G1 (*solid lines*) and S phase (*dashed lines*) cells in the experiment presented in panel C. Values represent data from a single experiment. See also Fig. S4.

We next tested the importance of the ΔΨm for normal cell cycle progression. To this end, we treated logarithmically growing cells with BAM15, an uncoupler that efficiently dissipates the ΔΨm without depolarizing the plasma membrane (Kenwood et al., 2014). Interestingly, BAM15 treatment of ρ^+^ cells caused a transient accumulation of cells in G1 phase (Fig. 4B, Fig. S4C). The observed accumulation of cells in G1 phase was accompanied by a corresponding decrease in the percentage of cells in S phase. Treatment with another uncoupler, CCCP, caused a similar transient increase in G1 phase cells as BAM15 (Fig. S4D-E).

To further corroborate the connection between ΔΨm and G1-to-S progression, we synchronized cells in G1 with α-factor and released them into medium containing BAM15. In analogy to the G1-to-S transition delay in ρ^0^ cells, BAM15 treatment delayed the progression of α-factor-synchronized ρ^+^ cells from G1 into S phase: while the percentage of G1 cells began to decline 20 min after release of ρ^+^ cells into normal medium, indicating transition from G1 into S phase, ρ^+^ cells released into BAM15-containing medium did not show signs of G1 exit until 70 min after release (Fig. 4C-D). BAM15-treatment also delayed the G1-to-S transition in ρ^+^ cells released from G2 synchrony: while the presence of BAM15 did not appreciably affect the transition from G2 to G1 phase, ρ^+^ cells released from G2 into BAM15-containing medium showed an increased percentage of G1 cells and delayed S phase entry, consistent with a G1-to-S transition delay (Fig. S4F-I).

The results of Figure 4 demonstrate that while the inhibition of mitochondrial ATP synthesis has no apparent impact on cell cycle progression, a ΔΨm collapse induced by uncoupler treatment causes cells to accumulate in G1 phase by triggering a G1-to-S phase transition delay similar to the one observed in ρ^0^ cells. These findings implicate ΔΨm in modulating cell cycle progression ρ^+^ cells.

### The cell cycle defect in ρ^0^ cells is exacerbated by a further reduction of membrane potential

In contrast to uncoupler-treated cells where the ΔΨm is largely dissipated, ρ^0^ cells have the ability to maintain a membrane potential that, although lower than in ρ^+^ cells, is still sufficient to support various ΔΨm-dependent processes like the import of nuclear-encoded mitochondrial proteins (Miceli et al., 2012; Appleby et al., 1999). Next, we examined whether the decreased ΔΨm could underlie the cell cycle phenotype of ρ^0^ cells and act as the initial signal to trigger the G1-to-S transition delay. We measured the ΔΨm of WT ρ^+^ and ρ^0^ cells using the fluorescent cationic dye tetramethylrhodamine methyl ester perchlorate (TMRE), and corrected the signal for potential changes in mitochondrial mass as detected by nonylacridine orange (NAO), a green-fluorescent dye that localizes to the mitochondria in a ΔΨm-independent manner (Doherty and Perl, 2017). Uncoupler-treated samples analyzed in parallel provided a measure of the background fluorescence. As expected, the ΔΨm of WT ρ^0^ cells was significantly lower than that in WT ρ^+^ cells, but clearly higher than the baseline value of uncoupler-treated cells (Fig. 5A, Fig. S6A-B).

**Fig. 5.**
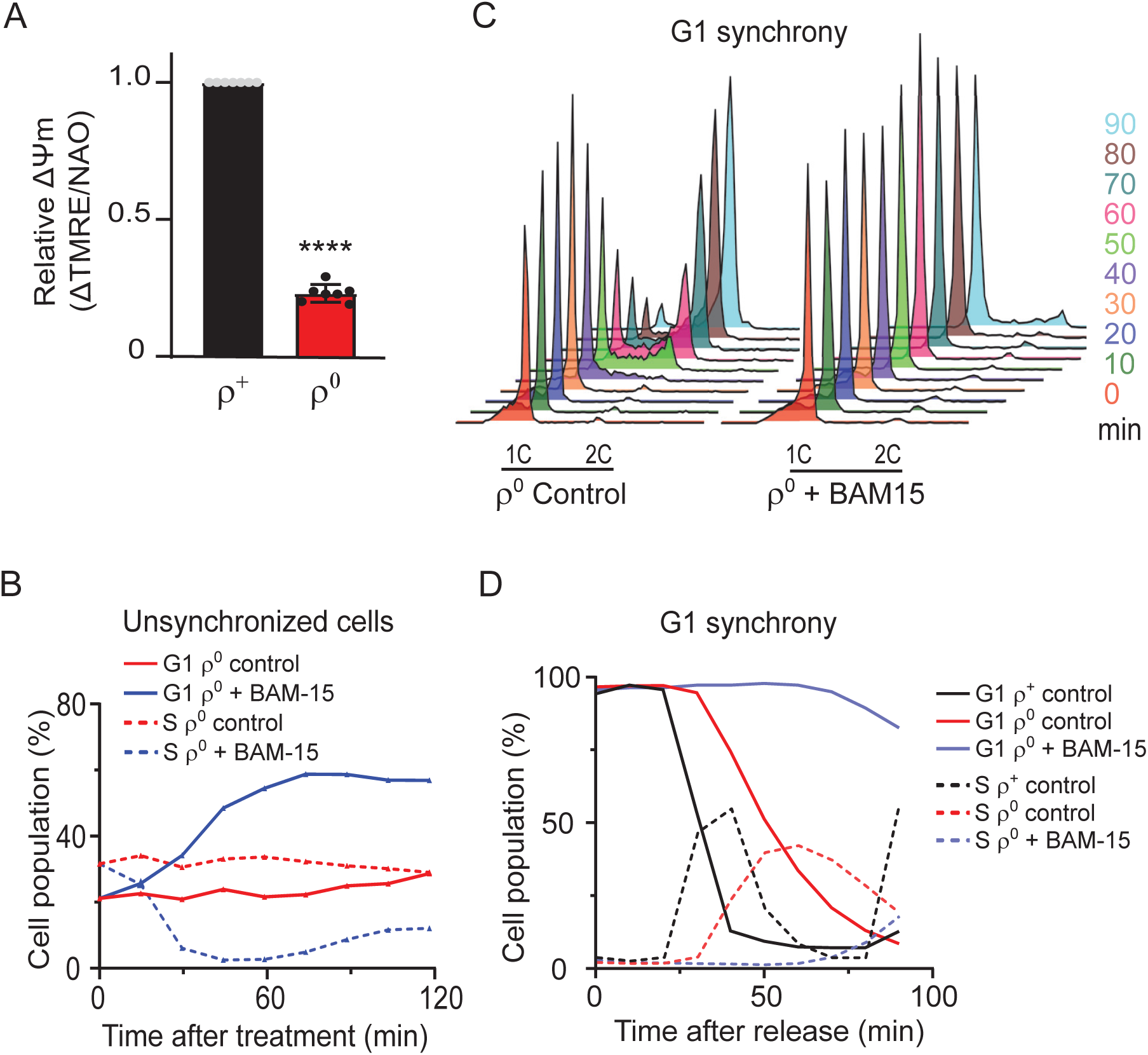
Uncoupler treatment exacerbates the cell cycle phenotype of ρ^0^ cells. **(A)** ΔΨm normalized to mitochondrial mass was measured in WT (AC403) ρ^+^ and ρ^0^ cells as described in Materials and Methods. The average of seven independent experiments is shown; error bars represent standard deviation. The groups were compared using one-way ANOVA; ****p <0.0001. The same data is shown in Fig. 6A. **(B)** The percentage of G1 (*solid lines*) and S phase (*dashed lines*) cells in cultures of WT ρ^0^ cells left untreated or treated with 30 μM of BAM15. Aliquots were harvested upon addition of the drug (0 min) and every 15 min thereafter. Values represent data from a single experiment repeated at least three times. Representative DNA histograms are shown in Fig. S5A. **(C)** DNA histograms of WT (AC402) ρ^0^ cells synchronized in G1 with 10 μg/ml α-factor and released into media with or without 30 μM BAM15. Aliquots were sampled upon release (time 0) and every 10 minutes thereafter. **(D)** Quantification of G1 (*solid lines*) and S phase (*dashed lines*) cells in the experiment presented in Fig. 5C (untreated and BAM15-treated ρ^0^ cells); the ρ^+^ control data is from Fig. 4C. Representative data from a single experiment is shown. See also Fig. S5.

In additional support of a role for ΔΨm in regulating cell cycle progression, the accumulation of ρ^0^ cells in G1 phase was further exacerbated when ΔΨm was fully dissipated by BAM15 or CCCP treatment (Fig. 5B, Fig. S5A-C). Furthermore, the effect of BAM15 was more sustained in ρ^0^ than in ρ^+^ cells and was evident even 2 h after addition of the compound (compare Fig. 5B and Fig. 4B). Expectedly, time course experiments following release from α-factor or nocodazole-induced synchrony revealed a more severe G1-to-S phase progression delay in ρ^0^ cells in the presence of BAM15 (G1 exit starting 40 min and 80 min after release from α-factor synchrony into medium lacking or containing BAM15, respectively) (Fig. 5C-D, Fig. S5D-F). These observations establish a quantitative correlation between ΔΨm and cell cycle progression, where the extent of the G1-to-S delay is governed by the severity of ΔΨm loss.

### *ATP1-111* rescues the cell cycle phenotype of ρ^0^ cells

P^0^ cells differ significantly from ρ^+^ cells in terms of metabolism given that respiratory-deficient ρ^0^ cells must compensate for the loss of a subset of the citric acid cycle reactions, the products of which are central to many anabolic pathways (Epstein et al., 2001). If the G1-to-S phase progression delay is indeed driven by decreased ΔΨm and not by other functional or metabolic differences between ρ^+^ and ρ^0^ cells, we hypothesized that it should be rescued by increasing ΔΨm in ρ^0^ cells. In the absence of ETC activity, mtDNA-deficient cells maintain ΔΨm by an alternative mechanism that involves the hydrolysis of glycolytically-produced ATP by the “reverse” action of the F1 subunit of the mitochondrial ATP synthase and the electrogenic exchange of ATP^4-^ for ADP^3-^ over the inner mitochondrial membrane (Appleby et al., 1999; Giraud and Velours, 1997). The *ATP1-111* mutation in the Atp1 subunit of the mitochondrial F1-ATPase results in a hyperactive enzyme that generates a higher ΔΨm in ρ^0^ cells than the one maintained in WT ρ^0^ cells (Francis et al., 2007; Veatch et al., 2009; Miceli et al., 2012). Accordingly, analysis of *ATP1-111* ρ^0^ cells revealed a ΔΨm comparable to WT ρ^+^ cells, confirming the *ATP1-111* ρ^0^ strain as a suitable model to test our hypothesis in (Fig. 6A, Fig. S6A-B).

**Fig. 6.**
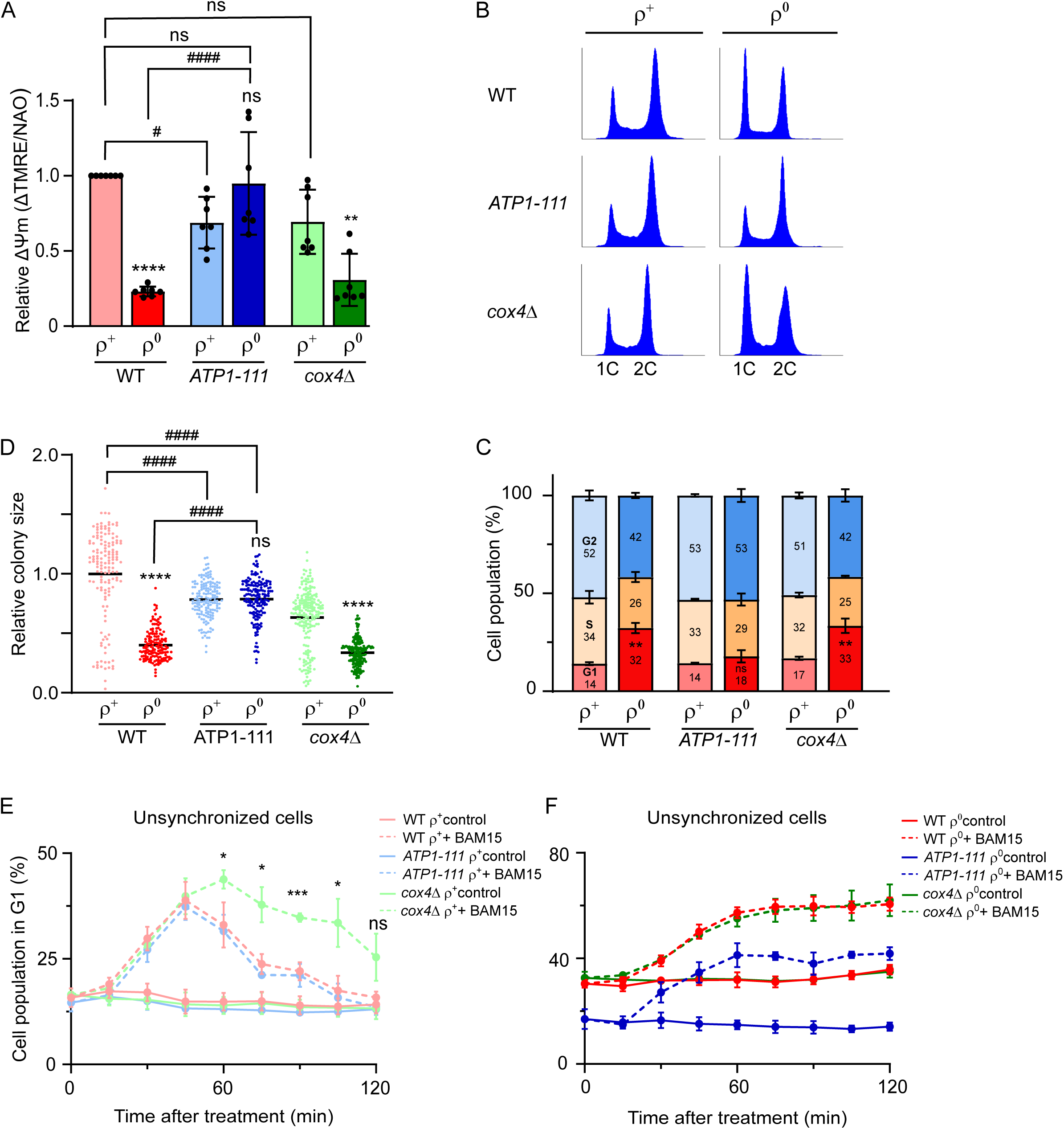
The cell cycle delay can be rescued by increasing the ΔΨm of ρ^0^ cells. **(A)** ΔΨm normalized to mitochondrial mass was measured in WT (AC403), *ATP1-111* and *cox4*Δ ρ^+^ and ρ^0^ cells as described in Materials and Methods. Histograms of TMRE and NAO fluorescence are presented in Fig. S6A-B. The average of seven independent experiments is shown; error bars represent standard deviation. The groups were compared by one-way ANOVA. ****p<0.0001, **p<0.01, *p<0.05, ns p>0.05 compared to respective ρ^+^; ####p<0.0001, #p<0.01 compared to indicated strain. WT ρ^0^ did not significantly differ from *cox4*Δ ρ^0^; WT ρ^+^ *vs. cox4*Δ ρ^0^: ####; WT ρ^0^ *vs. ATP1-111* ρ^+^: ##. **(B)** Representative DNA histograms of WT (AC403), *ATP1-111* and *cox4*Δ ρ^+^ and ρ^0^ cells grown to early-logarithmic phase in YPDA. **(C)** Quantification of the percentage of cells in G1, S or G2 phase in panel B. Values represent the average of at least 3 independent experiments, and error bars indicate standard deviation. The two-tailed Student’s t-test was performed to determine statistical significance between the G1 populations of each strains ρ^+^ and ρ^0^ variants. **p <0.01, ns p>0.05. **(D)** Quantification of the colony sizes of WT (AC403), *ATP1-111* and *cox4*Δ ρ^+^ and ρ^0^ cells grown on the same YPDA plate for 48 h as shown in Fig. S6C; the average area of WT ρ^+^ colonies was set to 1. The groups were compared by one-way ANOVA. ****p<0.0001, ns p>0.05 compared to the respective ρ^+^; ####p<0.0001 compared to the indicated strain. **(E-F)** The percentage of WT (AC403), *ATP1-111* and *cox4*Δ ρ^+^ (**E**) and ρ^0^ cells (**F**) in G1 phase in untreated cultures (*solid lines*) or after treament with 20 μM of BAM15 (*dashed lines*). The average of three independent experiments is shown; error bars represent standard deviation. Percentage of cells in G1 phase between WT and *cox4*Δ ρ^+^ cells at specific time points was compared by two-tailed Student’s t-test.***p<0.001, *p<0.05, ns p>0.05. See also Fig. S6.

We first analyzed the cell cycle profile of unsynchronized WT and *ATP1-111* ρ^+^ and ρ^0^ cells. As expected based on the relatively high ΔΨm in *ATP1-111* ρ^+^ cells, the strain showed a cell cycle profile that was indistinguishable from WT ρ^+^ cells (Fig. 6B-C). Remarkably, analysis of *ATP1-111* ρ^0^ cells revealed that the *ATP1-111* mutation was able to rescue the cell cycle phenotype of the WT ρ^0^ cell: while WT ρ^0^ cultures contained 32% of G1 phase cells, in *ATP1-111* ρ^0^ cultures the percentage of G1 cells was only 18% and thus comparable to the value in *ATP1-111* ρ^+^ cells (14%). Moreover, the colony size of *ATP1-111* ρ^0^ did not differ significantly from *ATP1-111* ρ^+^ cells (Fig. 6D, Fig. S6C), and their growth rate was partially restored relative to WT ρ^0^ cells (Fig. S6D) as previously reported by others (Francis et al., 2007; Veatch et al., 2009).

The earlier experiments carried out on WT cells revealed a more sustained effect of BAM15 on the cell cycle profile of ρ^0^ than ρ^+^ cells (Fig. 4B and Fig. 5B). We attributed this to the slower recovery of ΔΨm in ρ^0^ cells that lack a functional ETC. To explore this aspect further, we monitored cell cycle progression in ρ^+^ cells lacking Cox4, a nuclear-encoded subunit of complex IV. Unperturbed *cox4*Δ ρ^+^ cells were able to maintain a ΔΨm that did not significantly differ from that of WT ρ^+^ cells (Fig. 6A, Fig. S6A-B). Accordingly, the deletion of the *COX4* gene did not have any obvious effect on the cell cycle profile of unsynchronized ρ^+^ cells (Fig. 6B). However, *cox4*Δ ρ^+^ cells exhibited a stronger and more sustained cell cycle response to BAM15 than WT or *ATP1-111* ρ^+^ cells, indicating that a functional ETC aids in timely recovery from uncoupling (Fig. 6E, Fig. S6E). In line with this notion, all three ρ^0^ strains, including the *ATP1-111* ρ^0^ with the restored ΔΨm, failed to recover from the BAM15-induced accumulation of G1 cells within the 120 min follow-up period, although the percentage of G1 cells was lower in *ATP1-111* ρ^0^ than in WT or *cox4*Δ ρ^0^ cells (Fig. 6F, Fig. S6E). As expected based on their comparable ΔΨm values, the cell cycle profile of *cox4*Δ ρ^0^ cells was similar to WT ρ^0^ cells (Fig. 6A; B-C; F).

Based on the findings in Figure 6, we conclude that the delayed G1-to-S phase progression in ρ^0^ cells can be attributed to a decreased ΔΨm. Moreover, the cell cycle defect and the consequent petite phenotype and slow growth of ρ^0^ cells can be rescued by increasing ΔΨm through the *ATP1-111* mutation, implicating membrane potential— rather than other metabolic adaptations in ρ^0^ cells — as the main determinant of the cell cycle delay. Accordingly, recovery from transient uncoupler-induced cell cycle delay is expedited by a functional ETC.

### Altered ROS is not the signal for the G1-to-S phase progression delay in ρ^0^ cells

Mitochondria are a major source of reactive oxygen species, and mitochondrial dysfunction is often associated with increased levels of intracellular ROS (Leadsham et al., 2013). Furthermore, ROS are documented to influence cell proliferation (Nunnari and Suomalainen, 2012). Therefore, we wanted to determine whether ROS levels play a role in the ρ^0^ G1-to-S phase progression delay. Rather than increasing the percentage of cells in G1 however, elevated ROS following treatment with H_2_O_2_ primarily caused accumulation of WT ρ^+^ cells in early S phase, as previously reported by others (Fig. 7A-B; Fig. S7A) (Leroy et al., 2001). In contrast, decreasing ROS by treatment of WT ρ^+^ cells with the antioxidants N-acetylcysteine (NAC) or reduced glutathione (GSH) caused a transient accumulation of cells in G1 phase along with a decrease in the percentage of cells in S phase (Fig. 7C upper panel; Fig. S7B). A similar but more sustained effect of treatment with either NAC or GSH was observed in WT ρ^0^ cells (Fig. 7C lower panel; Fig. S7B). On their own, these data would suggest that neutralizing ROS by antioxidant treatment inhibits G1-to-S transition. However, measurement of ΔΨm in antioxidant-treated ρ^+^ cells revealed considerable effects of the antioxidants on ΔΨm, with a decrease of comparable extent to that of uncoupler treatment (Fig. 7D; Fig. S7C). Similar effects of NAC on ΔΨm have been reported in mammalian cells (Jiao et al., 2016; Al-Nahdi et al., 2018). The impact of antioxidant treatment on *S. cerevisiae* cell cycle progression observed in Fig. 7C may therefore be indirect and attributable to decreased ΔΨm rather than lower ROS levels *per se*.

**Fig. 7.**
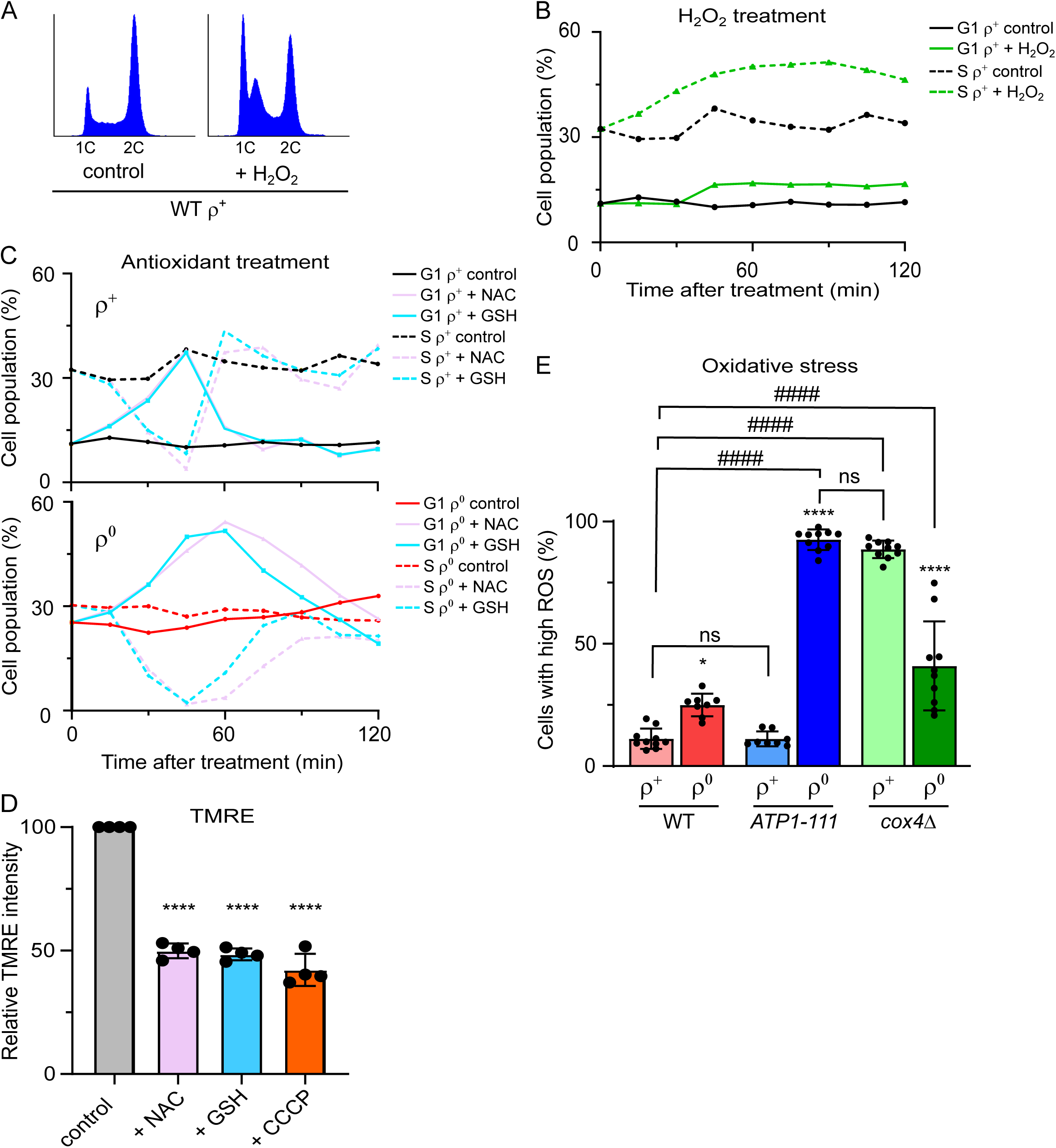
Levels of oxidative stress do not correlate with the extent of the G1-to-S delay. **(A)** A representative DNA histogram of WT (AC402) ρ^+^ cells grown until early logarithmic stage and treated with 400 μM H_2_O_2_ for 60 mins. Additional timepoints are shown in Fig. S7A. **(B)** Quantification of G1 (*solid lines*) and S phase (*dashed lines*) cells in the experiment presented in Fig. 7A. **(C)** Quantification of % of G1 (*solid lines*) and S phase (*dashed lines*) cells in WT ρ^+^ (upper panel) and WT ρ^0^ (lower panel) strains treated with 30 mM NAC or 20 mM GSH for 2 h. DNA histograms are shown in Fig. S7B. **(D)** Quantification of the TMRE uptake of WT ρ^+^ cells treated with 30 mM NAC, 10 mM GSH or 30 μM CCCP. Relative TMRE uptake was calculated using the geometric mean of the flourescence intensities of 20,000 cells, and the value of the untreated control was set to 100. The average of four independent experiments is shown; error bars represent standard deviation. The control and the treated samples were compared by one-way ANOVA. ****p <0.0001. A histogram from a representative experiment is shown in Fig. S7C. **(E)** Quantification of the percentage of cells with high levels of oxidative stress as measured by the fluorescence intensity of H_2_DCFDA. The average of ≥8 independent experiments is shown; error bars represent standard deviation. Comparison between variants and strains was performed by one-way ANOVA. ****p<0.0001,*p<0.05, ns p>0.05 compared to the corresponding ρ^+^; ####p<0.0001 or ns p>0.05 compared to the indicated strain. A representative experiment is shown in Fig. S7D. See also Fig. S7.

To further explore the relationship between ROS, ΔΨm and cell cycle progression, we measured the levels of oxidative stress in our strains of interest using the fluorescent probe H_2_DCFDA that reacts with several forms of ROS including H_2_O_2_, hydroxyl and peroxyl radicals (Jakubowski and Bartosz, 2000). As expected (Leadsham et al., 2013), the percentage of cells with an H_2_DCFDA signal above baseline levels (“high-ROS cells”) was two-fold higher in WT ρ^0^ cells than in WT ρ^+^ cells (Fig. 7E, Fig. S7D). Even *ATP1-111* ρ^0^ cells exhibited a higher percentage of high-ROS cells than the corresponding ρ^+^ strain. Interestingly, while *ATP1-111* ρ^+^ cells showed a comparable percentage of cells with high ROS as WT (11-12%), the percentage of high-ROS cells in *ATP1-111* ρ^0^ was as high as 92% and thus *ca* 8-fold higher than in WT or *ATP1-111* ρ^+^ cells. Therefore, the rescued cell cycle phenotype in *ATP1-111* ρ^0^ cells cannot be attributed to decreased ROS. In line with a lack of correlation between ROS and the cell cycle delay in ρ^0^ cells, in the *cox4*Δ pair of strains, mtDNA-devoid cells had a lower rather than a higher percentage of high-ROS cells when compared to the corresponding ρ^+^ cells.

Taken together, these data indicate that although neutralizing intracellular ROS by antioxidant treatment inhibits G1-to-S transition, this effect is likely indirect and mediated by diminished ΔΨm. In support of this interpretation, ROS levels did not correlate with timely cell cycle progression in the yeast strains analyzed. Rather, the collective findings of this study uncover ΔΨm as a novel modulator of cell cycle progression in eukaryotic cells.

## Discussion

The complexes of the ETC and ATP synthase are encoded on both cellular genomes, making retrograde communication from the mitochondria to the nucleus a prerequisite for mitochondrial biogenesis and the correct assembly of these critical complexes. Mito-cellular signaling is also required to trigger any compensatory responses that allow the cell to adjust to mitochondrial dysfunction. Therefore, inter-organellar communication initiated in the mitochondria is crucial under both physiological and pathological conditions.

A number of studies have implicated signals of mitochondrial status in the control of cell cycle progression. Mitochondrial dysfunction brought about by ETC mutations was shown to delay G1-to-S transition in *Drosophila melanogaster* imaginal disc cells (Mandal et al., 2005; Owusu-Ansah et al., 2008). Interestingly, these studies found different mitochondrial cues to be responsible for triggering the G1/S cell cycle checkpoint depending on the underlying ETC subunit mutation: while a complex IV mutation that decreased ATP levels signaled through AMPK and p53 to prevent S phase entry, a complex I mutation induced high ROS and signaled through the Foxo/p27 pathway. These findings highlight the fact that the signals and pathways that modulate the cell cycle in response to mitochondrial dysfunction can vary even within a single type of cell.

In *S. cerevisiae*, the identity of the mitochondrial signal that triggers the accumulation of respiratory-deficient ρ^0^ cells in G1, a response known as the mtDNA inheritance checkpoint, has remained unknown. In the current study, we rule out the two cues of mitochondrial dysfunction identified in *Drosophila*—loss of mitochondrial ATP production and high ROS—as triggers of the yeast cell cycle delay. Instead, we show that the G1-to-S transition delay observed in ρ^0^ cells is caused by decreased ΔΨm. Accordingly, the cell cycle phenotype of ρ^0^ cells could be recovered by increasing ΔΨm, indicating that low ΔΨm rather than respiratory deficiency *per se* restricts cell cycle progression in ρ^0^ cells (Fig. 5-6). Dissipation of ΔΨm in ρ^+^ cells induced a similar but transient G1-to-S delay (Fig. 4). These results therefore establish mitochondrial membrane potential as a general modulator of cell cycle progression in *S. cerevisiae* cells.

In mammalian cells, the ΔΨm has been reported to fluctuate over the course of the cell cycle, with the highest ΔΨm, mitochondrial O_2_ consumption and ATP synthesis measured just prior to the G1-to-S transition (Schieke et al., 2008; Mitra et al., 2009). Furthermore, Mitra *et al*. elegantly demonstrated that the boost in ΔΨm in late-G1 cells was required for S phase entry (Mitra et al., 2009). The ΔΨm therefore appears to regulate cell cycle progression even in higher eukaryotes. In addition, the mitochondrial membrane potential is known to govern *e*.*g*. mitochondrial dynamics and quality control through mitophagy (Meeusen and Nunnari, 2005; Jin et al., 2010), highlighting its role as a central readout of mitochondrial status.

In several models of mitochondrial dysfunction, the G1-to-S progression defect is dependent on the p53 checkpoint protein (Mitra et al., 2009; Mandal et al., 2005). Given that baker’s yeast lacks a p53 homolog, we explored the dependence of the ρ^0^ cell cycle phenotype on the upstream central checkpoint kinases Tel1, Mec1 and Rad53. Along with the RTG retrograde signaling pathway that is activated upon diminished ΔΨm (Miceli et al., 2012), these checkpoint proteins were found to be redundant for this phenomenon, ruling out both the RTG and the canonical checkpoint pathways as mediators of the ρ^0^ cell cycle delay (Fig. 2-3). The identity of the signaling pathway(s) that connect loss of ΔΨm to the cell cycle machinery therefore remains unclear. Because decreased ΔΨm can impair transport across the mitochondrial inner membrane, we speculate that the signaling mechanism may involve alterations in *e*.*g*. Ca^2+^ balance and/or protein import. Another interesting question is whether other energy metabolic readouts such as ROS or the levels of key metabolites can modulate the ΔΨm-dependent effects on the cell cycle. These and many related aspects of the mito-cellular communication that regulates the cell cycle progression of ρ^0^ cells should be addressed in future work.

Mitochondrial dysfunction underlies a diverse group of diseases that is caused by defects in mitochondrially-localized proteins encoded on either the nuclear or the mitochondrial genome. The common feature of mitochondrial diseases is defective function of the respiratory chain and/or ATP synthase that leads to insufficient mitochondrial ATP synthesis. In most cases, mitochondrial disorders also involve decreased ΔΨm (Chung et al., 2021b). However, not all symptoms of mitochondrial disease can be directly attributed to the shortage of ATP, as some stem from (mal)adaptive responses to the mitochondrial dysfunction. For example, dysregulated immune signaling triggered by mtDNA instability was shown to aggravate the metabolic dysfunction in a mouse model with an increased mtDNA mutation load (Lei et al., 2021). Similarly, patient fibroblasts carrying the common m.3242 A>G mtDNA point mutation exhibited constitutive activation of the PI3K-Akt-mTORC1 signaling axis, the inhibition of which partly improved mitochondrial function (Chung et al., 2021a). These examples illustrate the importance of understanding the various forms of mito-cellular crosstalk and the possible therapeutic potential of inhibiting signaling that in some cases may be more detrimental than beneficial. It remains to be seen whether inhibition of normal cell cycle progression in response to decreased ΔΨm contributes to any of the symptoms of mitochondrial disease.

## Materials and Methods

### Yeast strains and growth conditions

Unless otherwise indicated, all *Saccharomyces cerevisiae* strains used in this study are congenic to W4069-4c which is in a W303 background (Chabes et al., 2003), and are listed in Table 1. To ensure fitness before each experiment, all strains were streaked out from glycerol stocks onto YPDA solid medium (1% yeast extract, 2% peptone, 50 mg/ L adenine, 2% glucose, and 2% agar). Unless otherwise indicated, yeast cells were grown in YPDA at 30°C with shaking at 180 rpm.

**Table 1.**
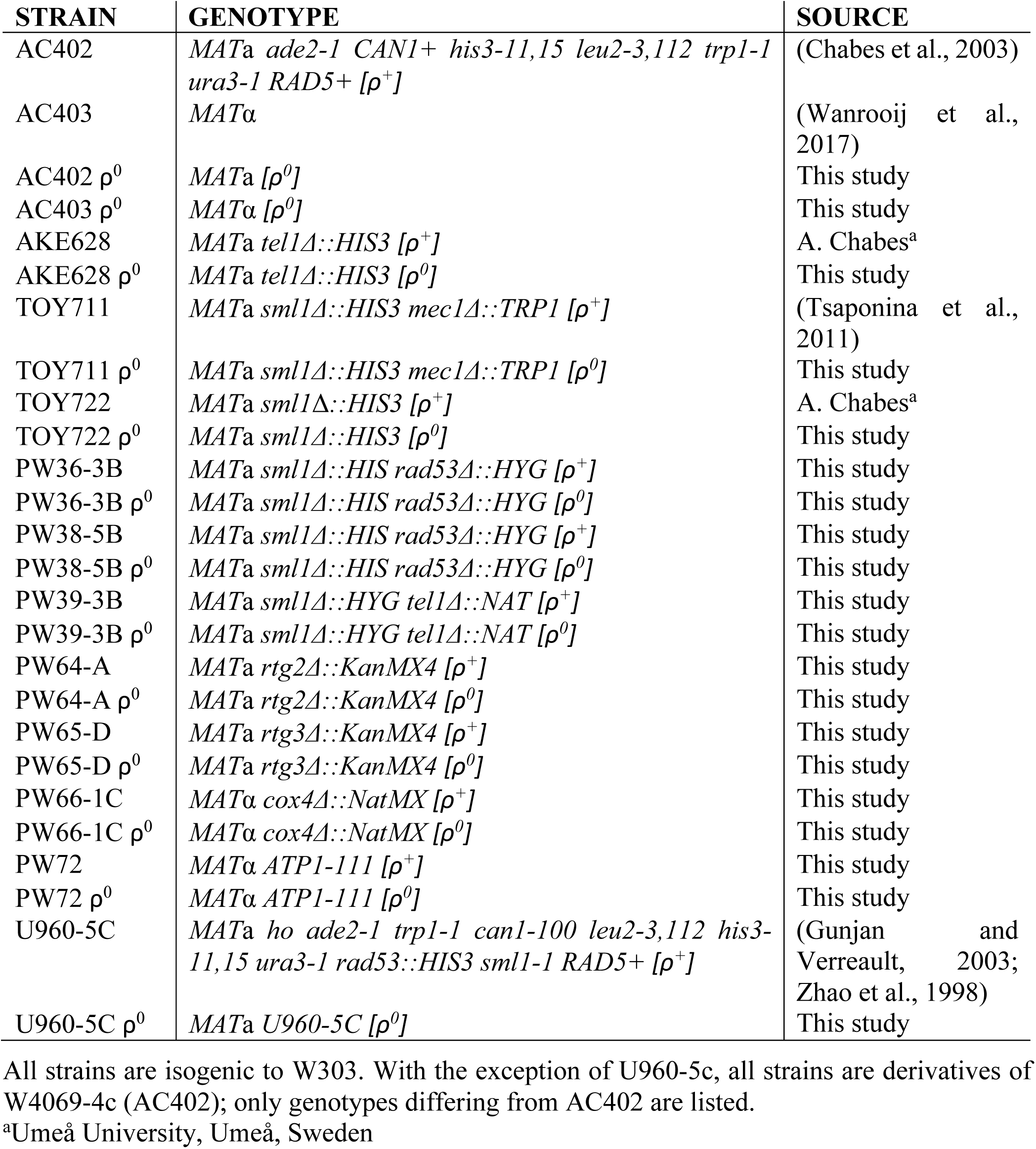
Yeast strains used in this study.

P^0^ strains lacking mtDNA were generated by growth in YPDA medium supplemented with 25 μM ethidium bromide for 4-15 days depending on the strain (Goldring et al., 1970). Loss of mtDNA was confirmed by lack of growth on YPGA medium (1% yeast extract, 2% peptone, 50 mg/ L adenine and 3% glycerol) in combination with the absence of amplification of a region of the mitochondrially-encoded *COX1* gene by quantitative real-time PCR in samples where a region of the nuclear-encoded *ACT1* gene amplified. Real-time PCR reactions contained 0.2 μM forward and reverse primers targeting *COX1* or *ACT1* (Table 2) and 10 μl 2x SyGreen Mix (PCRBiosystems, USA), and were run on a LightCycler 96 instrument (Roche, Switzerland) using the following program: 95°C 180 sec, 45 cycles of (95°C 10 sec, 56°C 10 sec, 72°C 1 sec with signal acquisition), melting curve (95°C 5 sec, 65°C 60 sec, heating to 97°C at 1 sec with continuous signal acquisition).

**Table 2.**
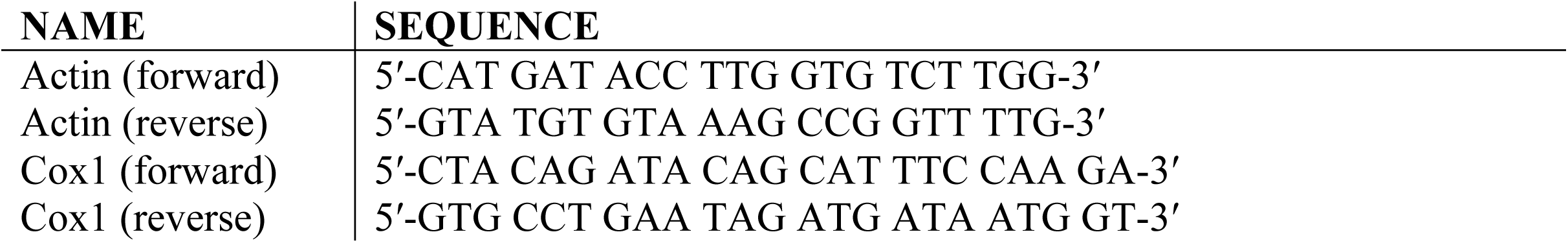
Oligonucleotides used in the study.

The strain bearing the *ATP1-111* mutation was generated by introduction of the *ATP1* variant carrying a T>G substitution at position 331 from a pRS316 integration vector as previously described (Francis et al., 2007; Miceli et al., 2012). Briefly, the *ATP1-111* insert in pRS306 was excised with BamHI and XbaI and re-cloned into the pRS316 integration vector. Yeast cells were transformed using a standard lithium acetate procedure (Gietz et al., 1995), and the presence of the mutation in cells that grew in medium supplemented with 5-FOA was confirmed by sequencing.

Deletion strains were constructed by replacing the entire open reading frame (ORF) with the *kanMX4* or *natMX* cassette (Bähler et al., 1998). Gene deletion was confirmed by growth in geneticin or nurseothricin, respectively, as well as by PCR using both a primer pair that flanks the ORF and a pair internal to it. PW36-3B, PW38-5B and PW39-3B were derived from backcrosses of TOY782 (Tsaponina et al., 2011), PY220 and PY304 (kind gifts of Dr. P. Burgers), respectively, to AC402/AC403.

### Colony size and growth curves

Yeast strains grown in YPDA liquid medium for at least 24 h were counted using a Neubauer Improved Hemocytometer and adjusted to 100 cells per 100 μl dH_2_O. 100 μl of the cell suspension was spread on a single partitioned YPDA plate and incubated at 30°C for 48 h prior to imaging on a ChemiDoc imaging system (Bio-Rad). The area of all colonies, except for ones with another colony attached to them, was analyzed using ImageJ and individual values were normalized to the average of all WT ρ^+^ cell colonies.

For growth curves, yeast strains cultured overnight in YPDA liquid media were re-inoculated in fresh media with starting OD_600_= 0.1 or 0.03. OD_600_ readings were recorded every 3 h by spectrophotometry.

### Cell cycle analysis by flow cytometry

Cells grown overnight in YPDA were diluted to a fresh medium to a starting OD_600_ of 0.1. Cells were harvested at OD_600_ 0.35-0.5, fixed with cold 70% ethanol and incubated at 4 °C overnight. The fixed cells were washed with distilled water once, resuspended in 50 mM Tris-HCl pH 7.5 with 15 mM NaCl, and treated with 2 mg/ml RNase A at 37°C overnight. The next day, samples were treated with 1.8 mg/ml proteinase K at 50 °C for 1 h, spun down and resuspended in 50 mM Tris pH 7.5. An aliquot was added to 50 mM Tris pH 7.5 containing a 1:10,000 dilution of SYBR green nucleic acid stain (Invitrogen, USA). The cell suspension was sonicated for 10 sec with an amplitude of 20% before analysis (QSonica Q500, USA). DNA content was detected in the FL1 channel using the Cytomics FC500 (Beckman Coultier, USA). Data analysis and quantification of cells in G1, S and G2 phases was performed using FCS Express 7 Flow (De Novo Software). The y-axis scale of the cell cycle histograms was adjusted to fit the highest cell count.

For time-course analysis of unsynchronized cells, 30 ml culture (with starting OD_600_ of 0.1) was grown to an OD_600_ of 0.4-0.6, split into two portions and the specified drug added to one portion while the other served as a control. Cell aliquots were harvested and immediately fixed every 15 minutes for 2 hours. Time 0 indicates addition of the drug.

For analysis of synchronized cells, cultures were synchronized in G1 by the addition of 5 μg/ml alpha-factor pheromone every hour for 2 h. For G2 synchronization, cells were treated with 10-25 μg/ml nocodazole for 2 hours. Cells were washed twice with one volume of ice-cold distilled water and released from synchrony by resuspension in fresh medium. Cells were harvested and immediately fixed every 10 min for 90-120 min following release.

### Mitochondrial membrane potential

Cells grown overnight in YPDA were diluted in fresh medium to a starting OD_600_ of 0.1 and grown for 24 h; 1 ml of the culture was harvested and washed twice with one volume of PBS buffer (0.14 M NaCl, 0.0027 M KCl, 0.010 M phosphate buffer pH 7.4). Cells were reconstituted in 5 ml PBS to a OD_600_ of 0.05. A 1 ml aliquot was treated with 2 μM tetramethylrhodamine methyl ester perchlorate (TMRE) (Molecular Probes, USA) and incubated at 37°C for 30 min. As control, another aliquot of the cell suspension was treated with the same concentration of TMRE for 20 min followed by addition of 20 μM carbonyl cyanide 3-chlorophenylhydrazone (CCCP) for 10 min to collapse the membrane potential. Mitochondrial mass was estimated by adding 250 nM nonylacridine orange (NAO) (Invitrogen, USA) to a third aliquot of the cell suspension and incubated at 37°C for 30 min. Both mitochondrial membrane potential and mitochondrial mass were determined by flow cytometry on a Beckman Coultier Cytomics FC500. ΔΨm was calculated by subtracting the TMRE fluorescence of the untreated and uncoupled sample and normalized for mitochondrial mass (Miceli et al., 2012).

### Measurement of cellular reactive oxygen species (ROS)

Determination of cellular ROS content was as previously described with some modifications (Gourlay and Ayscough, 2005; Yi et al., 2018). Briefly, yeast cells previously grown on solid medium were inoculated in YPDA and incubated at 30 °C for 24 hours. Cultures were diluted in 10 ml fresh medium to a starting OD_600_ of 0.1 followed by addition of 10 μg/ml H_2_DCFDA (Invitrogen, USA) and incubation at 30°C for 24 hours. One ml of the culture was harvested, washed twice with PBS and diluted to an OD_600_ of 0.1 with PBS. The cell suspension was sonicated for 10 sec with an amplitude of 20% (QSonica Q500, USA) before analysis by flow cytometry on a Beckman Coultier Cytomics FC500. Another set of cultures without H_2_DCFDA were analyzed to determine baseline fluorescence; the percentage of H_2_DCFDA-stained cells with higher than baseline fluorescence was used as a measure of oxidative stress (“high-ROS” cells).

### Western blot

Strains were grown in YPDA medium overnight, diluted to an OD_600_ of 0.1 the next morning and grown until OD_600_ 0.5-0.6. Cells were harvested by centrifugation, cell pellets flash-frozen in liquid nitrogen and stored at -80°C. As a positive control for Rad53 phosphorylation, DNA damage was induced where indicated by treatment with 2 μg/ml 4-Nitroquinoline 1-oxide (4-NQO) (Sigma) for 60 min before harvesting. Whole-cell protein extracts were prepared by bead-beating the pellets with 0.5 mm zirconia/silica beads (BioSpec) in 20% trichloroacetic acid (Scharlau) for 2 min. Protein extracts were pelleted and resuspended in 1 M Tris pH 7.6. One volume of 2 x Laemmli buffer (80 mM Tris, 8 mM EDTA, 120 mM DTT, 12% glycerol, 3.5% SDS) was added to the protein extracts and samples were incubated at 95°C for 5 min prior to loading onto a 10% polyacrylamide SDS-PAGE gel and resolving in Tris-Glycine-SDS buffer (25 mM Tris, 192 mM glycine, 0.1% SDS). Proteins were then transferred to a PVDF membrane (Amersham Hybond) in transfer buffer (25 mM Tris, 192 mM glycine, 20% ethanol). The membrane was blocked in TBS-T (20 mM Tris pH 7.6, 150 mM NaCl, 0.1% Tween-20) containing 1% BSA for 1 h at room temperature (RT), and incubated with primary antibodies against Rad53 (1:4000; rabbit, ab104332, Cell Signalling) or Tubulin (1:2000; rat, YL1/2, Abcam) diluted in TBS-T containing 1% BSA for 1 h 30 min at RT. After washing with TBS-T, the membrane was incubated with HRP-linked anti-rabbit (1:20 000; 31460, Thermo Scientific) or anti-rat (1:20 000; A5795, Sigma) secondary antibodies diluted in TBS-T with 1% BSA for 1 h at RT. After extensive washes with TBS-T, the blots were developed by chemiluminescence (ECL Bright, Agrisera) and images were taken using the ChemiDoc imaging system (Bio-Rad).

## Acknowledgements

We thank Drs. Andrei Chabes (Umeå University, Umeå, Sweden), Peter Burgers (Washington University in St. Louis, MO, USA) and Akash Gunjan (Florida State University, FL, USA) for yeast strains, and Drs. Michal Jazwinski (Tulane University, LA, USA) and Anders Byström (Umeå University, Umeå, Sweden) for plasmids. We acknowledge Dr. Sushma Sharma for expert technical advice and helpful discussions, and the P. Wanrooij laboratory for critical reading of the manuscript. This project was supported by grants from the Swedish Cancer Society (19 0022JIA and 19 0098Pj 01 H), the Swedish Research Council (2019-01874), the Swedish Society for Medical Research (S17-0023), the Kempe Foundations (JCK-1830) and the Åke Wiberg Foundation (M20-0132) to P.H.W. The authors declare no competing financial interests.

## Author contributions

C.M. Gorospe designed and carried out the majority of the experiments, analyzed the resulting data, supervised some activities, visualized all data and wrote parts of the manuscript. A.H. Curbelo, L. Marchhart and K. Niedźwiecka performed key experiments and analyzed the connected data. G. Carvalho conceptualized, designed and carried out some experiments, analyzed the connected data, contributed to supervision, and helped in writing the first draft. P.H. Wanrooij conceptualized the study, acquired funding, supervised the project, and wrote parts of the manuscript. All authors reviewed and edited the final version of the manuscript.

